# Integration of spatial single-cell proteomics and spatial metabolomics reveals tumor microenvironment predictive of immunotherapy response in mucosal melanoma

**DOI:** 10.64898/2026.04.23.720453

**Authors:** Jun Wang, Priyadharsini Nagarajan, Sungnam Cho, Yunhe Liu, Erin H. Seeley, Yibo Dai, Yang Liu, Kai Yu, Jared K. Burks, Jennifer L. McQuade, Adi Diab, Linghua Wang, Suhendan Ekmekcioglu

**Author notes:** Corresponding Authors: Dr. Suhendan Ekmekcioglu, Department of Melanoma Medical Oncology, Melanoma Medical Oncology, MD Anderson Cancer Center, University of Texas, 1515 Holcombe Blvd, Houston, TX, 77030; E-Mail; Dr. Linghua Wang, Department of Genomic Medicine, The University of Texas MD Anderson Cancer Center, 1881 East Road, 3SCR6.4111, Houston, TX, 77054. These authors contributed equally to this study (Jun Wang, Priyadharsini Nagarajan). These authors jointly supervised this study (Linghua Wang, Suhendan Ekmekcioglu).

## Abstract

Mucosal melanoma (MuM) is a rare but aggressive malignancy with limited benefit from immune checkpoint inhibition and few predictive biomarkers. We integrated single-cell spatial proteomics (COMET) and spatial metabolomics (MALDI-IMS) to profile 97 tissue cores from 26 patients treated with PD-1/PD-L1 and/or CTLA-4 inhibitors. We profiled 695,444 cells and resolved 25 cell states across eight major cell types. Cellular neighborhood (CN) analysis revealed distinct tumor- and stromal-associated spatial architectures. Responders were enriched for tumor-associated CNs (invasive tumor and tumor boundary) with close spatial proximity among Ki67⁺ tumor cells, CD163⁺ macrophages, and CD11c⁺ dendritic cells (DCs), and increased proliferating/cytotoxic CD8+ T-cell subsets. Non-responders showed stromal CN dominance with reduced immune infiltration. Spatial metabolomics identified lower abundance of indole-derived metabolites and reduced indole/tryptophan pathway activity in responders within tumor and TME regions that tracked with DC/macrophage-enriched spatial contexts. This study advances MuM spatial biology and provides a framework for biomarker-driven immunotherapy strategies.

**Statement of significance:** Mucosal melanoma (MuM) responds poorly to immune checkpoint blockade, and predictive biomarkers are limited. Integrated spatial proteomics and metabolomics reveal response-associated tumor-immune neighborhood architecture, stromal contexts linked to immune exclusion, and altered indole/tryptophan metabolism in the microenvironment. These spatial features nominate biomarkers and therapeutic hypotheses to improve immunotherapy for MuM.

## Introduction

Mucosal melanoma (MuM) is a rare and aggressive subtype of melanoma that arises from melanocytes located in mucosal tissues, including the nasal cavity, oropharynx, genitourinary tract, anorectum, and other mucosal sites (1). Despite accounting for only 1–2% of all melanomas, MuM is associated with a disproportionately high morbidity and mortality, with 5-year overall survival rates below 25% in most cohorts (2,3). Unlike its cutaneous counterpart, which is strongly associated with UV-induced mutagenesis, MuM typically displays a lower tumor mutational burden and a distinct genomic landscape, characterized by recurrent alterations in *KIT*, *NRAS*, and *BRAF* and *SF3B1*, together with frequent chromosomal alterations (4,5). Clinically, MuM often presents at advanced stages due to its occult anatomical location and lack of early symptoms, complicating timely diagnosis and effective intervention (6).

Recent advances in treatment with immune checkpoint inhibitors (ICIs) directed against PD-1/PD-L1 as monotherapy or combined with anti–CTLA-4 ICI have revolutionized the treatment landscape for cutaneous melanoma (7–9). In contrast, MuM frequently exhibits significantly lower response rates than in cutaneous disease, with reported objective response rates to anti–PD-1 monotherapy ranging from 15–25%, and slightly better responses with combination ICI therapy (10,11). In MuM, the genomic context—particularly alterations in *KIT* and *NRAS*—may influence response to ICI. Retrospective analyses suggest that patients harboring *KIT* mutations may derive comparable or slightly improved benefit from anti–PD-1 or anti–PD-L1 therapy compared to wild-type tumors, potentially reflecting differences in tumor-immune states, including enhanced immune cell infiltration and/or antigen presentation programs reported in *KIT*-altered disease (12). In contrast, *NRAS*-mutant MuM generally exhibit poorer responses to ICI. *NRAS* activation is linked to sustained MAPK signaling and has been reported to promote immunosuppressive cytokine programs, with diminished cytotoxic T-cell infiltration (13,14). However, mutation status alone offers an incomplete explanation, and robust predictive biomarkers that connect oncogenic programs to tumor-immune interactions and ICI outcomes in MuM remain lacking.

Emerging evidence further suggests that divergent responses to anti–PD-1 therapy among melanoma subtypes have been linked to distinct tumor immune profiles (15,16). These observations indicate that MuM harbors a tumor microenvironment (TME) that differs from cutaneous melanoma in ways that may favor immune evasion and limit therapeutic efficacy. Importantly, beyond cellular composition, the spatial architecture of the TME can strongly influence immune control and therapeutic response (17–19). The positioning of immune cells relative to tumor nests, as well as their interaction with stromal barriers, vascular compartments, and immunosuppressive niches, can shape whether cytotoxic lymphocytes access malignant cells or remain excluded at the periphery (17,18,20,21). To interrogate these spatial determinants in patient tissues, spatial proteomics has emerged as a transformative approach for high-dimensional, spatially resolved profiling of protein expression in situ (22). By quantifying immune and stromal markers across tumor regions while preserving tissue context, spatial proteomics provides high-resolution maps of cellular phenotypes, signaling pathways, and their microenvironmental context—offering insight into immune exclusion, tumor–stroma interactions, and candidate resistance mechanisms (23).

In parallel, spatial metabolomics has emerged as a complementary approach that maps the distribution of small-molecule metabolites directly within intact tissue sections (24,25). Unlike conventional bulk metabolomics, spatial metabolomics preserves native tissue architecture, enabling regional metabolic programs to be linked to local immune infiltrates, stromal compartments, and tumor regions or histologic subtypes within the same specimen (26,27). This spatially resolved metabolic layer adds functional context beyond proteomics and can help pinpoint metabolic pathways associated with immune suppression or activation (28). While the immune architecture of cutaneous melanoma and other solid tumors has been studied using high-dimensional spatial technologies, comparable analyses in MuM remain limited (29,30). As a result, it is still unclear how spatial heterogeneity, including immune cell exclusion and stromal compartmentalization shapes tumor-immune interactions and contributes to differential responses to ICI therapy in MuM.

To address this knowledge gap, we implemented a spatially resolved multi-omics approach combining COMET—a high-plex imaging platform capable of profiling 40 protein markers simultaneously—with matrix-assisted laser desorption/ionization imaging mass spectrometry (MALDI-IMS) to map and quantify 632 metabolite features. Using these complementary modalities, we analyzed 97 tissue cores from 26 patients with MuM, including both pre-treatment and post-treatment specimens. We then applied computational frameworks for cellular neighborhood inference, spatial context modeling, and cell–cell proximity/interaction profiling to define spatial programs linking immune infiltration, stromal compartmentalization, and regional metabolic states. Together, these analyses identified spatial features—spanning protein-defined cellular phenotypes and localized metabolic signatures—associated with immune activity, stromal organization, and clinical response to ICI therapy.

By integrating spatial single-cell proteomics and spatial metabolomics, our study provides a comprehensive, spatially resolved view of microenvironmental composition and architecture associated with response and resistance to immune checkpoint blockade in MuM. This integrative framework enables us to delineate patterns of tumor–immune organization, characterize stromal contexts linked to immune exclusion, and uncover metabolite programs linked to therapeutic outcomes. Together, these analyses establish a spatial multi-omic reference for immune dynamics in MuM and motivates spatially informed hypotheses and therapeutic strategies for this highly aggressive melanoma subtype.

## Results

### Spatial single-cell phenotyping of MuM using spatial proteomics

To profile the spatial heterogeneity of MuM, we implemented a multi-omic workflow integrating spatial proteomics (COMET) with spatial metabolomics (MALDI-IMS) (**Fig. 1A**). We developed a 40-antibody COMET panel (**Fig. 1B, left**), enabling multiplexed detection of a wide range of features, including MuM tumor markers, markers of major immune subsets (e.g., CD4⁺ and CD8⁺ T cells, NK cells, macrophages, B cells, and dendritic cells), and stromal compartments (**Fig. 1B, left**). This panel also included markers reporting functional states, including proliferation, antigen presentation, and inflammatory signaling (**Fig. 1B, left**). In parallel, MALDI-IMS was performed to map 632 metabolite features across tissue sections. Together, these modalities were applied to 97 formalin-fixed paraffin-embedded (FFPE) tissue cores from 26 patients.

**Figure 1.**
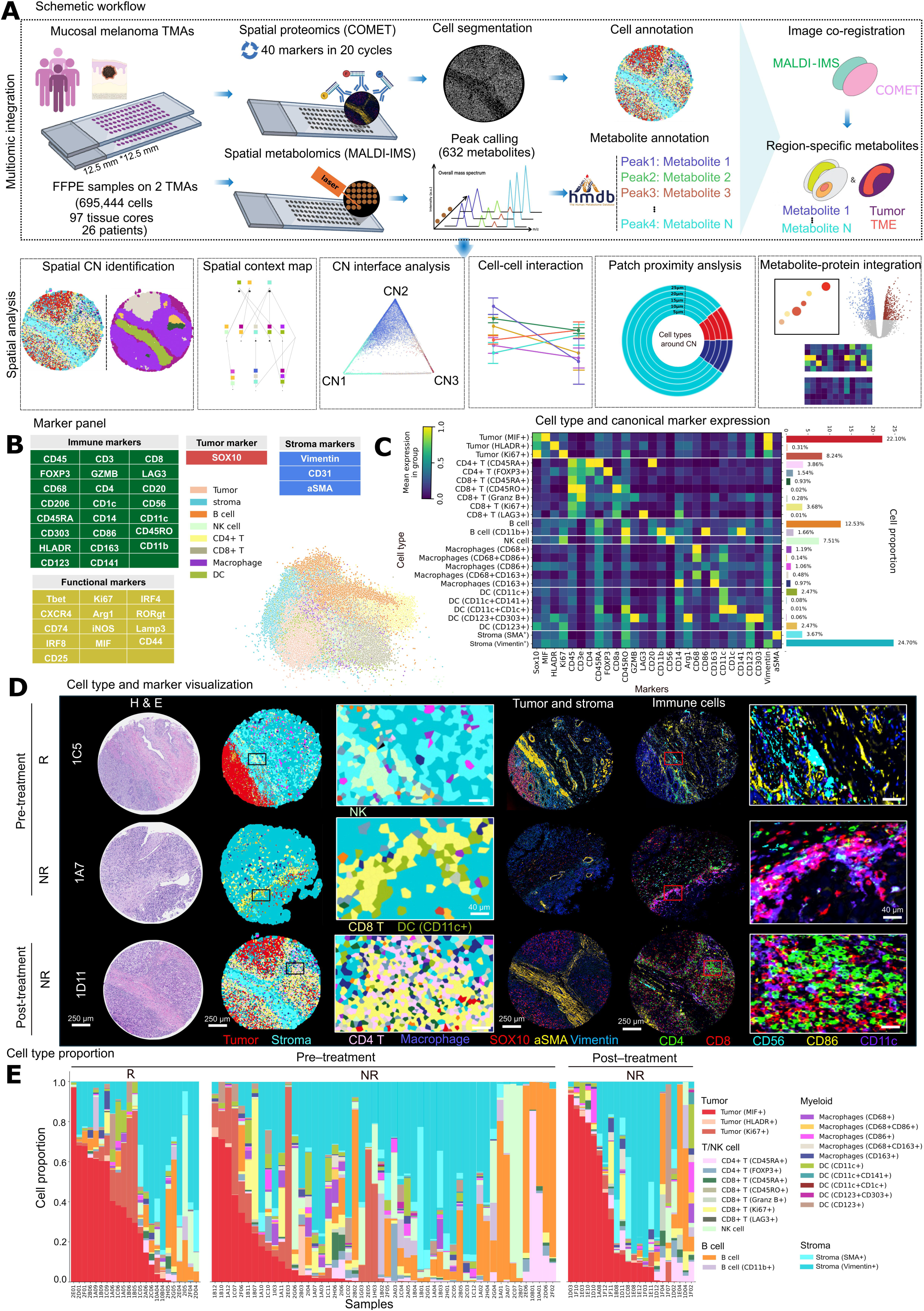
Study design and definition of cell types and phenotypes. (A) Overview of the study workflow integrating single-cell spatial proteomics (COMET) and spatial metabolomics (MALDI-IMS) for profiling MuM tissue cores. Consecutive tissue sections were subjected to 40-plex COMET imaging and MALDI-IMS, followed by multi-omics integration, region annotation, and downstream statistical and spatial analyses. (B) Antibody marker panel used for COMET profiling. Partial Least Squares Discriminant Analysis (PLS-DA) of a representative subset of single cells, colored by major cell lineages defined by marker expression profiles listed at left. (C) Heatmap showing normalized mean marker expression across the 25 identified cellar phenotypes. Bar plots (right) indicate the relative abundance of each phenotype across all tissue cores. (D) Representative tissue regions shown with H&E staining (left), Voronoi plots annotated by cell type (middle), and COMET images colored by cell type-specific marker expression (right). (E) Stacked bar plots illustrating the relative proportions of all cell types across all tissue cores, stratified by pre-treatment R, pre-treatment NR and post-treatment NR cores.

Following stringent quality control, we retained a total of 695,444 segmented single cells for downstream analysis. Unsupervised clustering and marker-based annotation identified 8 major cell types (**Fig. 1B, right**), which were further resolved into 25 cellular phenotypes (**Fig. 1C**). We visualized cell segmentation and phenotype assignments using Voronoi tessellation plots from representative specimens, including pre- and post-treatment cores from a non-responder (NR) and a pre-treatment core from a responder (R) (**Fig. 1D**). We quantified cell state compositions for each sample and compared pre- and post-treatment timepoints across response groups (i.e., pre-treatment R, pre-treatment NR, and post-treatment NR). Overall, we observed marked differences in the composition of immune and stromal phenotypes and their spatial arrangement across these groups (**Fig. 1E**). Specifically, immune cells were detected across tumor regions, whereas infiltration levels varied substantially across cores and patients, reflecting both intratumoral and interpatient heterogeneity.

We next summarized cell-type composition across samples. Overall, the proportions of major TME lineages were not significantly different between pre-treatment R and NR cores (**Supp. Fig. 1A and 1B)**, and between pre- and post-treatment NR cores (**Supp. Fig. 1C and 1D**. In contrast, examining tumor phenotypic states revealed differences: MIF⁺ tumor cells were significantly more abundant in pre-treatment R cores and in post-treatment NR cores compared with pre-treatment NR cores (**Supp. Fig. 1E**). Conversely, Ki67⁺ tumor cells were significantly higher in NR pre-treatment cores than in NR post-treatment cores, and were also elevated in NR relative to R in the pre-treatment group (**Supp. Fig. 1E and 1F**). Together, these data indicate that while global TME composition alone may not distinguish response groups at baseline, specific tumor cell states differ by clinical response and treatment exposure.

### Distinct spatial cellular neighborhoods in MuM

To characterize spatial cellular architecture in MuM, we performed cellular neighborhood (CN) analysis, which quantifies local cellular composition using a window-based spatial model across the tissue as previous reported (31) (**Fig. 2A**). This approach identified 15 distinct CNs, which were characterized by their cellular composition (heatmap; **Fig. 2B**, left), their relative abundance across samples (**Fig. 2B**, right), and their spatial distribution within tissues (**Supp. Fig. 2**). Within tumor regions, four biologically distinct tumor CNs were identified (**Fig. 2B**): CN6 (proliferative tumor) was enriched for Ki67⁺ tumor cells, consistent with locally of proliferative areas (**Fig. 2C**, bottom left). CN15 (central tumor) was enriched for MIF⁺ tumor cells, a tumor state associated with immunomodulatory programs, and was predominantly localized to tumor cores (32) (**Fig. 2C**, top left). CN14 (invasive tumor) comprised a mixture of tumor cell states (Ki67⁺, MIF⁺, and HLA-DR⁺), consistent with a transitional interface with phenotypic diversity (**Fig. 2C**, top left). CN13 (tumor boundary) contained tumor cells together with infiltrating macrophages, consistent with a tumor-myeloid interface at the margin (**Fig. 2C**, top left). Of note, the central tumor CN (CN15) was surrounded by the invasive tumor CN (CN14). In turn, CN14 was bordered by the tumor boundary CN (CN13).

**Figure 2.**
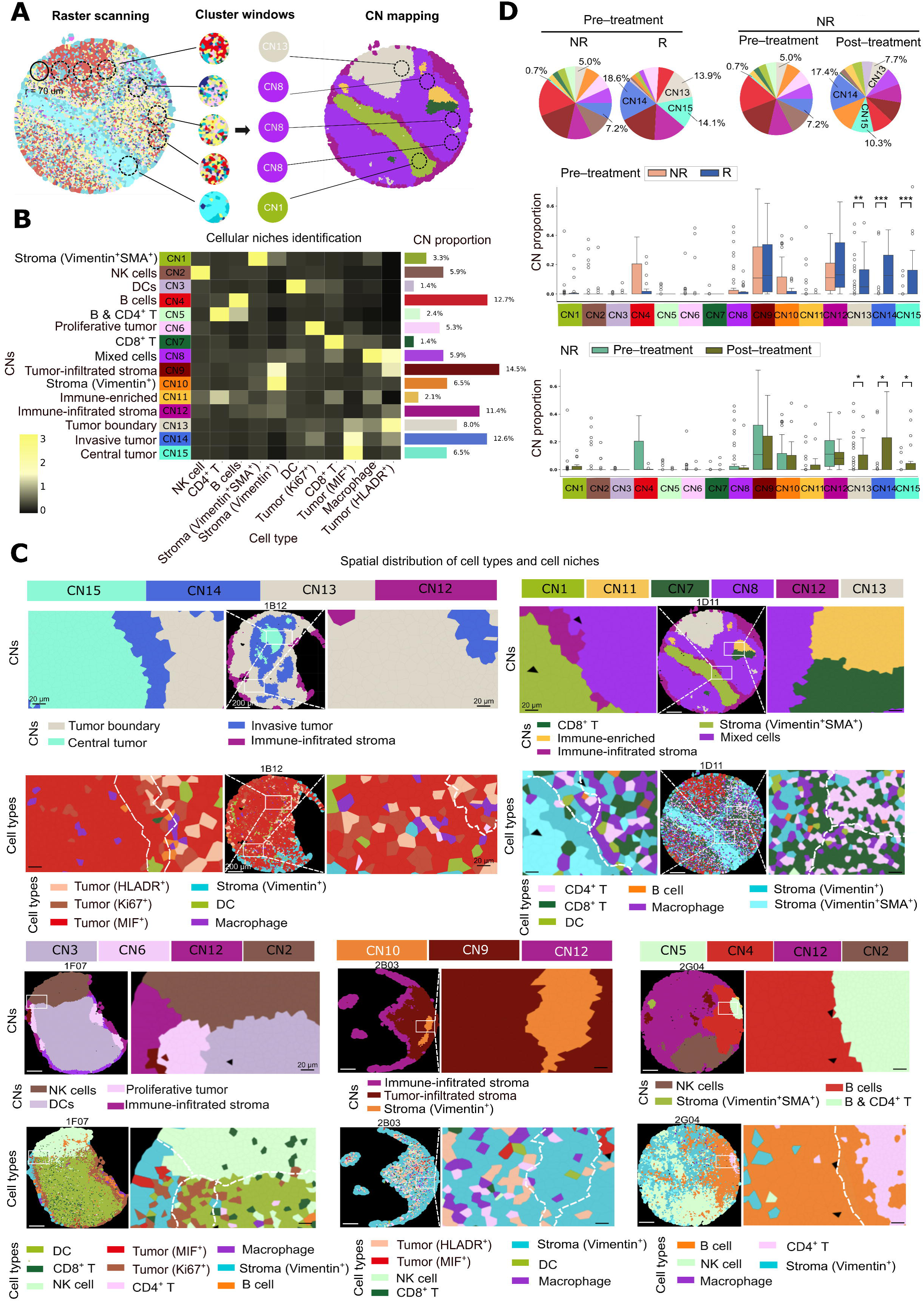
Identification and characterization of CNs in MuM. (A) Schematic illustration of CN identification and mapping (left), with color-coded CNs displayed across a representative tissue core (right). (B) Identification of 15 distinct CNs based on local cell composition. Left: overview of CN types; right: relative proportions of each CN across all tissue cores. (C) Voronoi diagrams colored by CNs (upper row) and cell types (lower row) within representative regions (left) and zoomed-in view (right) for each CN. (D) Boxplots comparing CN frequencies between pre-treatment R and pre-treatment NR cores, as well as between pre-treatment NR and post-treatment NR cores. Each dot represents the CN frequency from an individual tissue core (*p < 0.05, **p < 0.01, ***p < 0.001; Mann–Whitney U test).

In stromal and tumor-adjacent compartments, four predominant CNs were identified (**Fig. 2B**). CN10 (Vimentin^+^ stroma) was enriched for vimentin^+^ stromal cells (**Fig. 2C**, bottom middle), whereas CN1 (Vimentin^+^SMA^+^ stroma) was enriched for SMA⁺ stromal cells consistent with a myofibroblastic cancer-associated fibroblast (myCAF) context (**Fig. 2C**, top right). CN9 (tumor-infiltrated stroma) and CN12 (immune-infiltrated stroma) were characterized by mixed stromal–tumor or stromal–immune cell populations, consistent with a spatially interdigitated stromal architecture (**Fig. 2C**, bottom middle). In immune-enriched compartments, multiple CNs corresponding to distinct immune cell populations were identified (**Fig. 2B**). CN3 (DC) was enriched for dendritic cells, while CN2 (NK cells) was enriched for natural killer cells (**Fig. 2C**, bottom left). CN4 (B cells) and CN5 (B & CD4⁺ T) displayed B cell–dominant and mixed B–CD4⁺ T cells, respectively (**Fig. 2C**, bottom right). CN7 (CD8⁺ T) was characterized by CD8⁺ T cell enrichment, whereas CN11 (immune-enriched) comprised a heterogeneous immune population (**Fig. 2C**, top right). CN8 (mixed cells) displayed a diverse cellular composition without dominance of a single lineage (**Fig. 2C**, top right).

Comparative analysis of CN frequencies further revealed that CN15, CN14, and CN13 were significantly enriched in pre-treatment R cores and in post-treatment NR cores (**Fig. 2D**), suggesting that treatment exposure and/or response status is associated with distinct spatial architectures of tumor-associated neighborhoods.

### Stromal context is associated with immune exclusion and ICI resistance in MuM

To examine higher-level tissue organization, we generated spatial context maps summarizing CN-CN adjacency patterns for each response/timepoint group (**Fig. 3A**). Across groups, the interface among three tumor-associated CNs–central tumor (CN15), invasive tumor (CN14), and tumor boundary (CN13)–was increased in pre-treatment R cores compared with pre-treatment NR cores (**Fig. 3A**, top). Beyond tumor-tumor neighborhood interfaces, CN combinations involving dendritic cell (DC)-associated CN combinations were particularly enriched in the pre-treatment R cores and post-treatment NR cores (**Fig. 3A**, top and bottom). Representative spatial distributions of DC-associated CNs in a responder specimen were shown in **Fig. 3B**. In parallel, stromal CN combinations also differed across groups (**Fig. 3A**, middle). Notably, pre-treatment and post-treatment NR cores showed increased combinations involving vimentin^+^ stroma CN (CN10), tumor-infiltrated stroma CN (CN9), and immune**-**infiltrated stroma CN (CN12). Representative stromal architectures across groups were illustrated in **Fig. 3C**.

**Figure 3.**
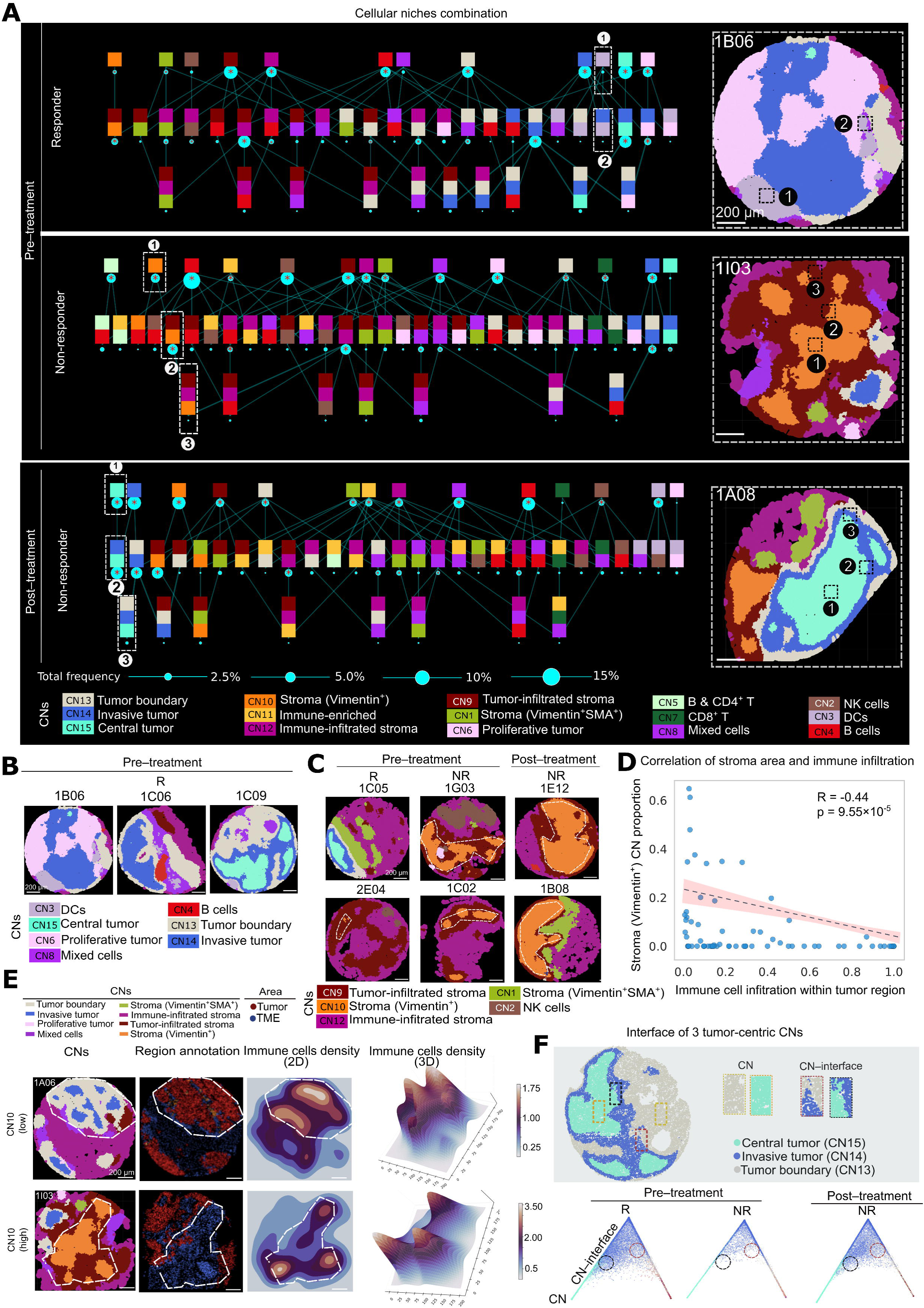
Spatial organization and inter-CN relationships associated with ICI response. (A) Left, spatial context maps summarizing hierarchical CN neighborhood combinations among pre-treatment R, pre-treatment NR and post-treatment NR tissue cores. Rows indicate the degree of neighborhood diversity: row 1 indicated dominance by a single CN (>85% composition), row 2 represented two dominant CNs (>85% combined), and subsequent rows denoted increasingly heterogeneous contexts. Right, representative tissue cores illustrating these multi-neighborhood configurations. (B) Representative spatial maps of DCs-enriched CN distribution in three pre-treatment R tissue cores. (C) Spatial maps illustrating stroma-associated CN combinations in representative pre-treatment R, pre-treatment NR and post-treatment NR tissue cores. (D) Correlation between stromal CN abundance and immune cell infiltration within tumor regions. (E) Representative spatial maps of CNs as well as tumor and stroma region annotations (left), with immune cell density visualized in two-dimensional and three-dimensional representations (middle and right). (F) Barycentric coordinate plots depicting the spatial interface among central tumor, invasive tumor, and tumor boundary CNs across pre-treatment R, pre-treatment NR and post-treatment NR tissue cores.

We next asked whether stromal architecture relates to immune access within tumor regions. Correlation analysis revealed a negative association between the abundance of CN10 and the extent of immune cell infiltration in tumor areas (**Fig. 3D**), suggesting that stromal components may form physical barriers limiting immune infiltration. This association was further supported by representative samples with high versus low CN10 abundance (**Fig. 3E**, left): corresponding 2D and 3D immune cell density plots consistently demonstrated that tumor regions with low CN10 harbored higher levels of immune infiltration, whereas samples with high CN10 showed reduced immune infiltration (**Fig. 3E**, middle and right).

### Spatial tumor-immune interactions are associated with ICI response

To further delineate immune architecture, barycentric coordinate mapping demonstrated that the interfaces between the invasive tumor CN (CN14) and central tumor CN (CN15) as well as invasive tumor CN (CN14) and tumor boundary CN (CN13) were enriched in pre-treatment R cores (**Fig. 3F**). Motivated by this observation, we next quantified cell-type composition across the three tumor-associated CNs—CN15, CN14, and CN13—and performed patch proximity analysis centered on CN13, examining its immune distributions within 25 μm (**Fig. 4A**). At the level of major immune lineages, pre-treatment R and NR cores showed broadly comparable distributions across tumor CNs and across the 5–25 μm boundary window (**Fig. 4B**, top). However, when focusing on immune subsets, distinct patterns emerged. In R cores, macrophages were more frequently of the CD163⁺ subtype and were more abundant than in NR cores. R cores also showed higher abundance of cycling CD8⁺ T (Ki67^+^) cells and cytotoxic CD8^+^ T (Granzyme B^+^) cells, whereas regulatory CD4⁺ T (FOXP3⁺) cells were less frequent than relative to NR cores (**Fig. 4B**, middle–bottom). Spatial mapping of a representative tissue core further corroborated these findings, demonstrating a clear spatial gradient in immune cell distribution, with enrichment of DCs, macrophages, CD4⁺ T and CD8⁺ T cells from the central tumor CN toward the tumor boundary CN, highlighting boundary-associated immune accumulation (**Fig. 4C**).

**Figure 4.**
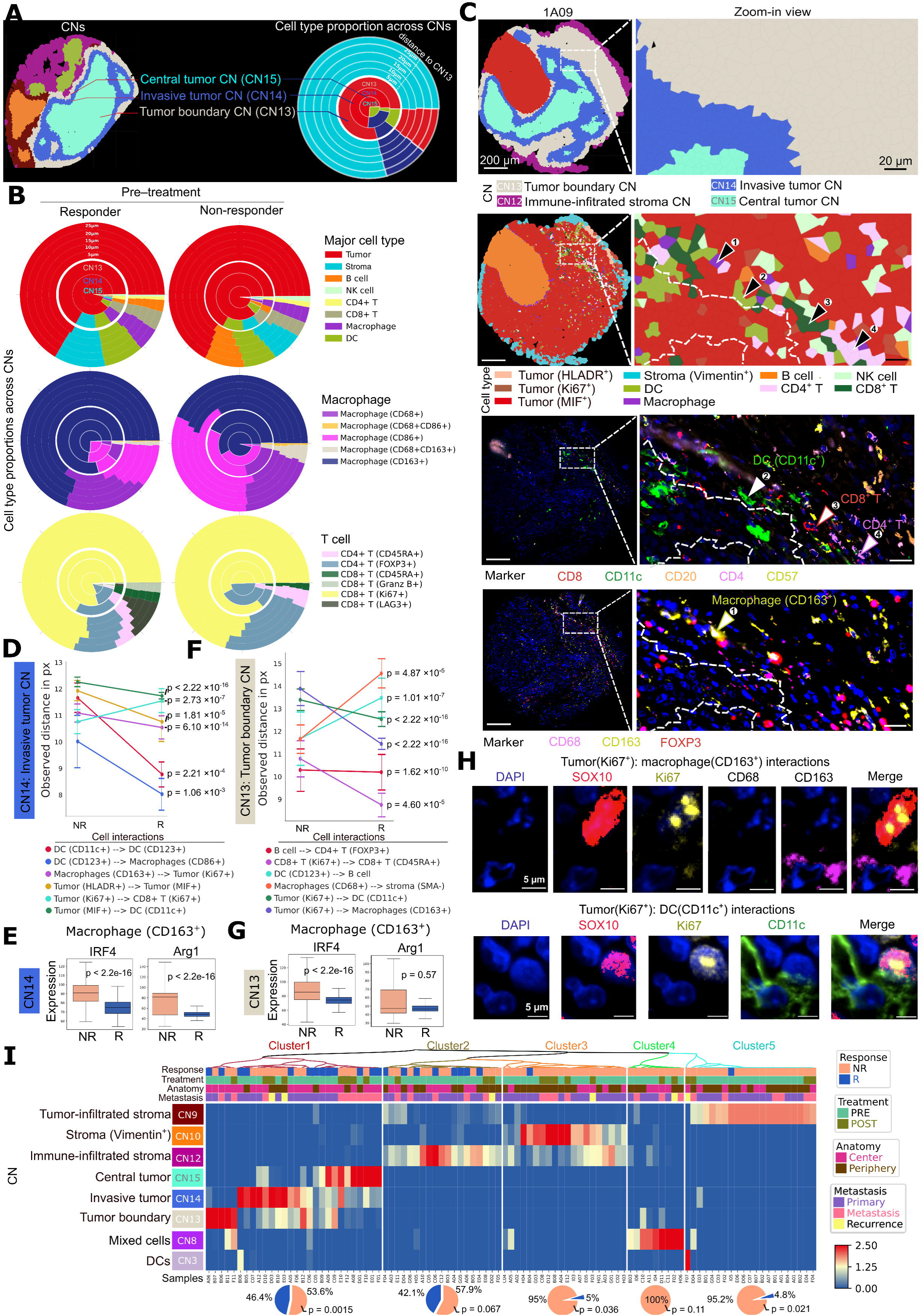
Spatial tumor–immune interactions associated with ICI response in MuM. (A) Schematic representation of tumor CNs (CN13, CN14 and CN15) (left) and spatial mapping of cell type proportions across eight layers, visualized as pie plots over a representative tissue core. (B) Relative abundance of major cell types (top), macrophage cell subtypes (middle) and T cell subtypes (bottom)— across tumor CNs and within 5–25 µm of patches from the tumor boundary CN in the pre-treatment R (left) and pre-treatment NR (right) cores. (C) Spatial mapping of tumor-associated CNs (CN13, CN14 and CN15), cell types, canonical markers and zoomed-in views of the selected region in one representative sample. (D) Top six statistically significant differential cell-cell interactions observed in the invasive tumor CN between pre-treatment R and pre-treatment NR cores. (E) Expression levels of key immunoregulatory markers (IRF4 and Arg1) in CD163⁺ macrophages within the invasive tumor CN, comparing pre-treatment R and NR cores. (F) Top six differential cell-cell interactions between pre-treatment R and NR cores within the tumor boundary CN. (G) Expression levels of IRF4 and Arg1 in CD163⁺ macrophage in the tumor boundary CN. (H) Representative high-resolution spatial images illustrating direct interactions between CD163⁺ macrophages, CD11c⁺ DCs and Ki67⁺ tumor cells. (I) Hierarchical clustering heatmap based on the z-scored frequency of cell phenotype-defined CNs across tissue cores.

R cores also exhibited differences in spatial proximity patterns within the tumor microenvironment. Specifically, R cores exhibited significantly closer spatial proximity between Ki67⁺ tumor cells and CD163⁺ macrophages in both the tumor boundary (CN13) (p < 2.22 × 10^−16^) and invasive tumor (CN14) (p = 6.10 × 10^−14^) CNs, and between Ki67⁺ tumor cells and CD11c⁺ DCs (p < 2.22 × 10^−16^) within the CN13 (**Fig. 4D** and **4F**). These findings were further illustrated by representative spatial images highlighting proximity between Ki67⁺ tumor cells and CD163⁺ macrophages (**Fig. 4H**, top) and between Ki67⁺ tumor cells and CD11c⁺ DCs (**Fig. 4H**, bottom). In contrast, CD163⁺ macrophages in NR cores expressed higher levels of IRF4 and Arg1 (**Fig. 4E** and **4G**), consistent with increased M2 polarization and a more immunosuppressive TME. Together, these observations suggest that a tumor-CD163⁺ macrophage/DC spatial community with reduced immunosuppressive features is associated with favorable response to ICI therapy in MuM.

Based on these findings, we next asked whether R and NR tissue cores could be stratified according to their CN profiles using an unsupervised approach. Hierarchical clustering of cell-phenotype–based CN compositions identified five distinct clusters (**Fig. 4I**). Cluster 1 comprised a mixture of R and NR cores (46.4% R and 53.6% NR) and was significantly enriched for R cores (p = 0.0015; pie plot, **Fig. 4I**). This cluster was characterized by enrichment of tumor-associated CNs, including CN15, CN14, and CN13 (heatmap; **Fig. 4I**). Cluster 2 consisted of 42.1% R and 57.9% NR cores and showed enrichment for R cores at trend level (p = 0.67; pie plot; **Fig. 4I**). Cluster 3 was predominantly composed of NR cores (95% NR and 5% R) and showed significant enrichment (p = 0.036; pie plot, **Fig. 4I**). Cluster 4 was composed exclusively of NR cores. Cluster 5 was also highly enriched for NR cores (95.2% NR and 4.8% R; p = 0.021; pie plot; **Fig. 4I**). Notably, Clusters 3 and 5 were more frequently associated with stroma-associated/infiltrated CNs (CN9 and CN10), in contrast to Cluster 1, which was dominated by tumor-associated CNs (heatmap; **Fig. 4I**). Together, these results support a model in which distinct spatial tumor–TME architectures are associated with differential responses to ICI therapy in MuM.

### Spatial metabolomics reveals altered fatty acids metabolism in MuM

To investigate regional metabolic differences between R and NR within tumor and TME compartments, we integrated spatial metabolomics using MALDI-IMS and identified 632 metabolite features across tissue sections (**Fig. 1**). We then selected the top 82 differentially abundant metabolites between R and NR for downstream analysis. Partial least squares discriminant analysis (PLS-DA) performed separately in tumor and TME compartments showed separation of R and NR based on these 82 metabolites (**Fig. 5A**). In the TME compartment, multiple unsaturated fatty acids (UFAs)-related metabolites—including linoleic acid, oleic acid, palmitate, and stearic acid—were specifically upregulated in pre-treatment samples of R compared with NR (**Fig. 5B**, right). Consistently, metabolite set enrichment analysis highlighted biosynthesis of UFAs remodeling as among the most significantly altered pathways within the TME compartment (**Fig. 5C**).

**Figure 5.**
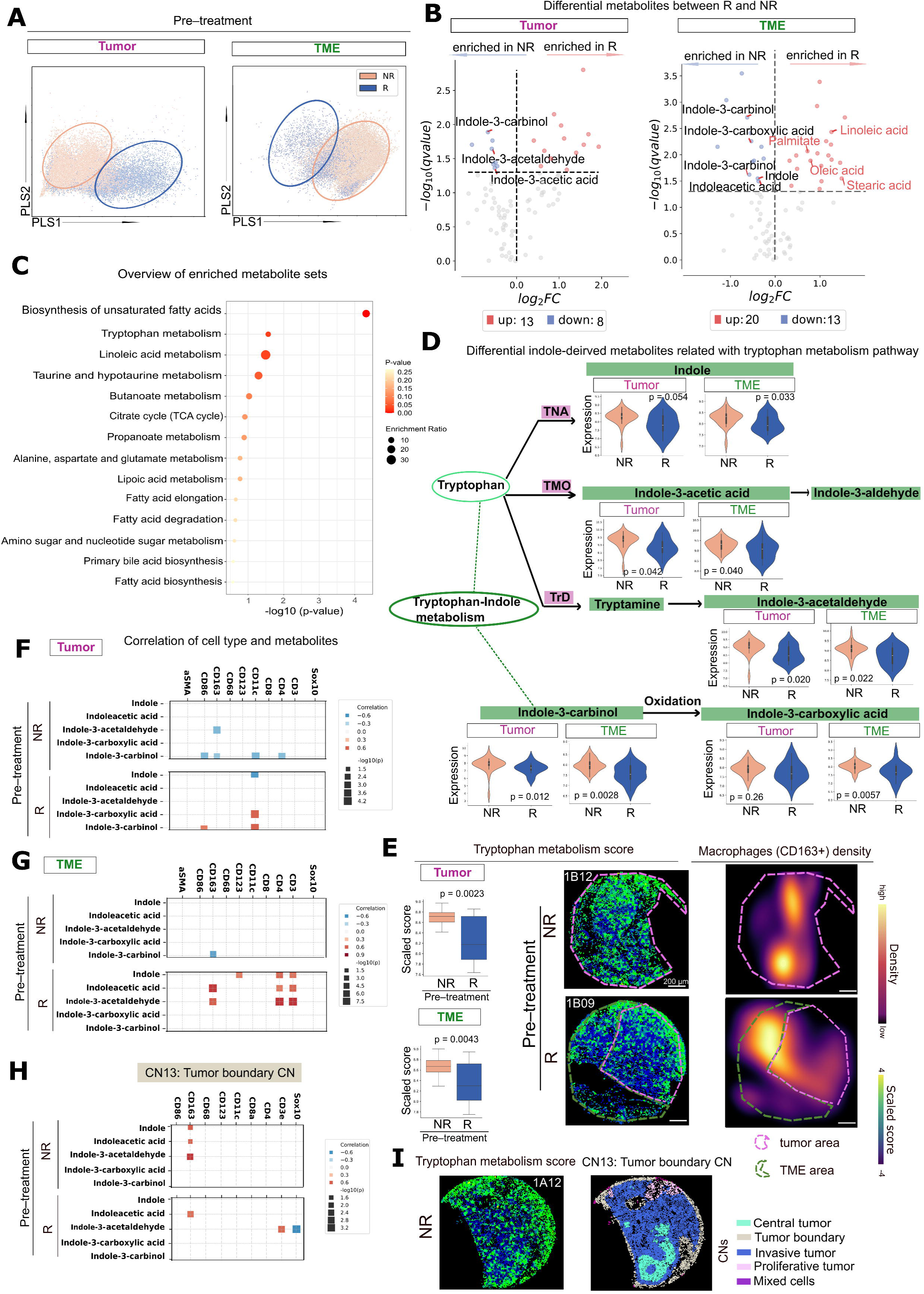
Spatial metabolomic alterations associate with ICI response in MuM. (A) PLS-DA analysis separating pre-treatment R and NR cores based on differentially abundant metabolites within tumor and TME compartments. (B) Volcano plot showing differential metabolite abundance between pre-treatment R and NR cores in the tumor and TME compartments. (C) Metabolite set enrichment analysis highlighting significantly altered metabolites in the TME compartment. (D) Schematic representation of tryptophan metabolism, with boxplots comparing the metabolites abundance between pre-treatment R and NR cores in tumor and TME compartments. (E) Tryptophan metabolism score comparing tumor and TME compartments between R and NR cores (left), and spatial distribution of two representative samples with CD163^+^ macrophages cell density (right). Correlation analysis between tryptophan metabolism–associated metabolites and canonical cell-type markers in the tumor compartment (F), TME compartment (G) and tumor boundary CN (CN13) (H). (I) Representative NR tissue core showing spatial distribution of tryptophan metabolism scores (left) and corresponding CN annotations (right). TNA: Tryptophanase; TMO: Tryptophan Monooxygenase; TrD: Tryptophan Decarboxylase.

### Tryptophan metabolism associates with ICI response in MuM

In pretreatment R cores, the tumor compartment exhibited reduced abundance of indole-3-carbinol and indole-3-acetaldehyde relative to NR cores (**Fig. 5B**, left). In the TME compartment, multiple indole-derived metabolites—including indole, indole-3-acetic acid, indole-3-carbinol, indole-3-acetaldehyde, and indole-3-carboxylic acid—were consistently downregulated in pre-treatment R cores compared with NR (**Fig. 5B**, right). Pathway enrichment analysis ranked tryptophan metabolism among the second most significantly altered pathways within the TME compartment (**Fig. 5C**). These findings were further illustrated in **Fig. 5D**, highlighting coordinated differences in metabolite abundance across the tryptophan metabolism pathway in both tumor and TME compartments. Consistent with these feature-level changes, a tryptophan metabolism score was reduced in R relative to NR in both tumor and TME compartments (**Fig. 5E**, left).

We next investigated metabolite patterns with spatial immune phenotypes. Indole-3-carboxylic acid and indole-3-carbinol were positively correlated with CD11c expression in the tumor compartment of pre-treatment R cores (**Fig. 5F**), consistent with spatial association between indole-pathway metabolites and CD11c⁺ DCs. In the TME compartment of pre-treatment R cores, indole-3-acetaldehyde and indole-3-acetic acid showed positive correlations with CD163 expression (**Fig. 5G)**, indicating that local indole-pathway abundance tracks with CD163⁺ macrophage-rich areas. In a representative pre-treatment R specimen, regions with high CD163⁺ macrophage density exhibited spatially higher tryptophan metabolism scores within the TME compartment (**Fig. 5E**, lower right), despite the overall reduction in tryptophan-pathway metabolites in R relative to NR at the group level. Similarly, in pre-treatment NR cores, indole, indole-3-acetaldehyde, and indole-3-acetic acid correlated positively with CD163 expression within the tumor boundary CN (CN13) (**Fig. 5H**), accompanied by elevated tryptophan metabolism scores in CN13 in NR patients (**Fig. 5I**), highlighting response-group differences in macrophage-associated immune–metabolite spatial coupling. Collectively, these integrated spatial proteomic and metabolomic analyses highlight a macrophage- and DC-associated tryptophan/indole metabolic axis that differs by response group and tissue compartment in MuM, suggesting spatial immune-metabolite interactions as potential contributors to variability in ICI response.

## Discussion

This study provides a high-resolution spatial and molecular characterization of MuM tumor cells and their surrounding TME, offering insights into cellular and metabolic features associated with response to ICI therapy. By integrating multiplexed spatial proteomics (COMET) with spatial metabolomics (MALDI-IMS), we delineated tumor-immune architecture, stromal contexts linked to immune exclusion, and metabolite signatures, particularly involving indole/tryptophan-related metabolites, that differed between R and NR cores. Key features distinguishing response groups are summarized in **Fig. 6**.

**Figure 6.**
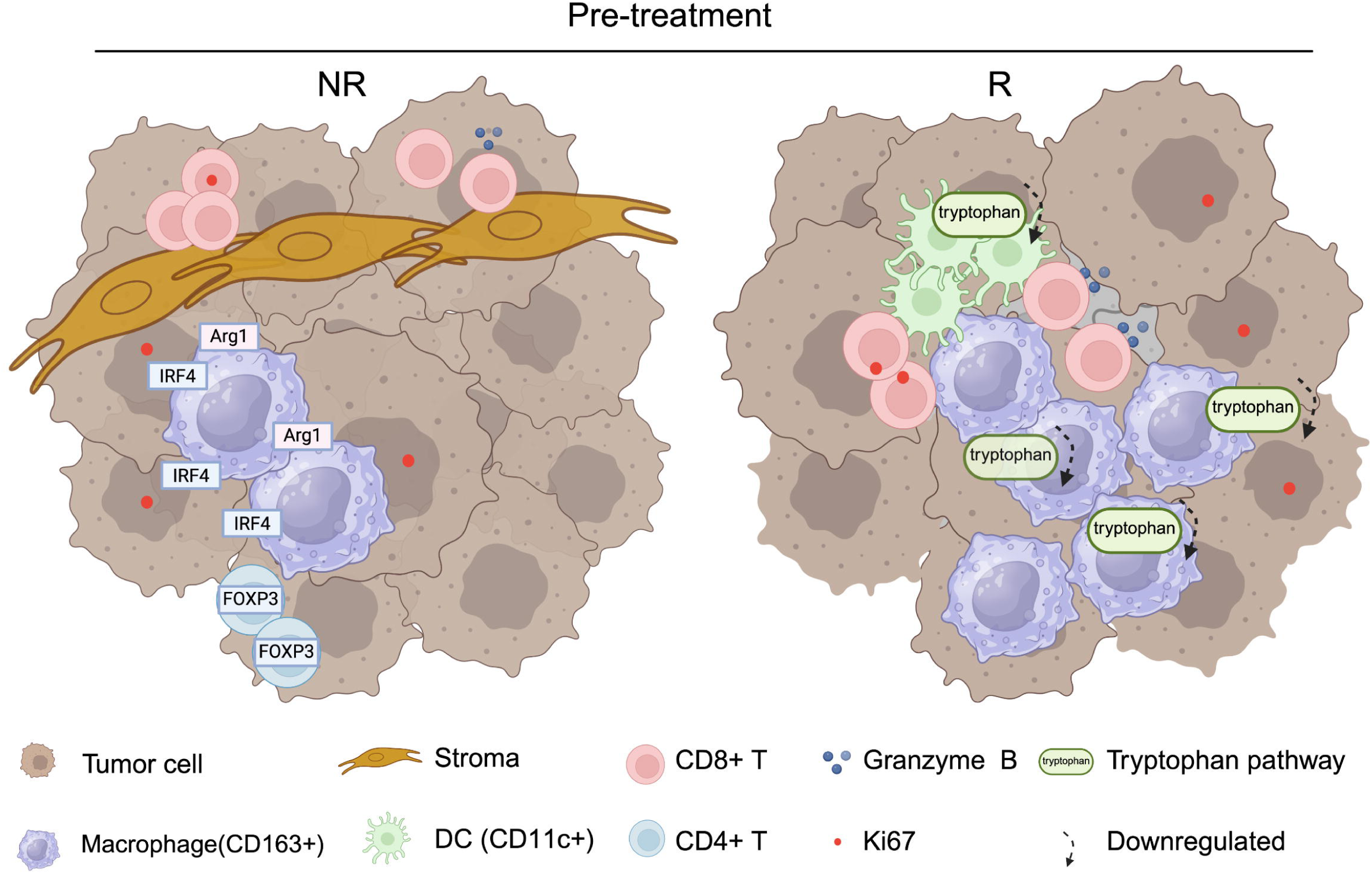
Summary of spatial immune and metabolic features distinguishing R and NR patients.

A central observation was the prominence of tumor-associated CN pattern, particularly the invasive tumor CN (CN14) and tumor boundary CN (CN13), in pre-treatment R cores. Within these tumor-associated CNs, R cores showed closer spatial proximity between proliferative tumor cells and CD163⁺ macrophages and CD11c⁺ DCs, together with increased frequencies of proliferating and cytotoxic CD8^+^ T cells. These spatial configurations are consistent with a TME that supports local antigen presentation, cytokine crosstalk, and efficient positioning of myeloid antigen-presenting cells near tumor regions–conditions that can favor productive T-cell priming and effector activity under ICI (33,34). Notably, CD163⁺ macrophages in R cores exhibited reduced expression of Arg1 and IRF4, consistent with attenuation of an immunosuppressive macrophage program and a shift toward a more immune-permissive state (35,36). While our data are correlative, they suggest that macrophage functional plasticity–and its spatial engagement with tumor cells and DCs–may be a key component of response-associated microenvironmental organization in MuM.

Together, these findings nominate myeloid context at the tumor interface as a potentially important determinant of response, where altered CD163⁺ macrophage program (e.g., reduced Arg1/IRF4) combined with proximity to tumor cells and DCs may shape antigen presentation and downstream T-cell activity under ICI. While causality remains to be established, macrophage reprogramming represents a plausible axis to explore for enhancing ICI therapy efficacy in MuM.

Beyond tumor-associated interfaces, DC-related CN combinations were more prominent in responders, supporting the concept that DC positioning and local context can be critical for initiating and sustaining antitumor T-cell responses (21), particularly in tumors with relative low baseline immunogenicity such as MuM (5,16). In line with this, we observed correlations between CD11c expression and specific indole-derived metabolites (e.g., indole-3-carboxylic acid and indole-3-carbinol), raising the possibility that local metabolic states may co-vary with DC-enriched microenvironments. These observations motivate future mechanistic studies to test whether metabolic cues directly influence DC activation (37), antigen presentation capacity, and downstream T-cell priming in MuM.

In contrast, the stromal compartment emerged as a major feature associated with reduced immune infiltration and therapeutic non-response. Non-responder tumors were significantly enriched for stromal CNs, and stromal abundance correlated inversely with immune cell infiltration. This supports the view that stroma may act as both physical and biochemical barriers, consistent with prior studies implicating extracellular matrix remodeling and CAF–mediated immunosuppression in impaired T cell trafficking (19,31,36,38,39). These findings underscore the potential biomarker value of stromal spatial features for immune exclusion and treatment failure, particularly within the constraints of core-based spatial sampling.

Our findings on metabolic profiling are significant in this rare melanoma subtype, with UFA-related metabolites among the most altered metabolic features. Altered lipid metabolism, particularly involving fatty acid remodeling and desaturation, has emerged as a hallmark of tumor progression and context-dependent immune regulation. Tumor cells actively remodel their lipid composition to support rapid proliferation, membrane biogenesis, and adaptation to metabolic stress. In this context, the desaturation of fatty acids—catalyzed primarily by stearoyl-CoA desaturase (SCD1)—plays a central role in maintaining membrane fluidity, redox balance, and signaling competence. Elevated SCD1 activity leads to increased synthesis of monounsaturated fatty acids (40,41). In colon cancer, high expression of SCD1 has been linked to reduced ICI sensitivity (42). Moreover, SCD1-mediated lipid remodeling has been shown to dampen T-cell activation and cytotoxicity (43). The role of UFAs in modulating immune cell behavior within the TME is also increasingly appreciated; Tumor-derived lipids can be taken up by macrophages and DCs, skewing them toward immunosuppressive phenotypes. In particular, exposure to unsaturated fatty acids can engage peroxisome proliferator–activated receptor (PPAR) signaling, promoting M2-like macrophage polarization and impairing DC cross-priming.

Notably, in our MuM cohort, selected UFAs were increased in R cores at the TME compartment, suggesting that the relationship between lipid states and ICI response maybe compartment- and context-dependent in MuM. While prior mechanistic work supports a role for lipid desaturation in immune resistance (42), our dataset does not directly establish tumor-intrinsic SCD1 activity as the driver of the observed metabolite differences. Overall, our study nominates spatial lipid metabolic heterogeneity as a response-associated feature and motivates future studies integrating lipid-pathway activity readouts (e.g., SDC1 expression/activity) and functional perturbations to evaluate whether targeting lipid remodeling can synergize with ICI in MuM.

Our metabolomic profiling analyses uncovered another significant alteration which is the suppression of tryptophan metabolism–derived indole compounds in R cores across tumor and TME regions. Particularly, indole-3-acetaldehyde and indole-3-acetic acid correlated with CD163 expression in tumor boundary and TME regions in pre-treatment R cores, potentially linking local macrophage-rich areas to spatial variation in indole/tryptophan pathway features (44). Given the immunosuppressive roles of tryptophan catabolites via activation of the aryl hydrocarbon receptor (AhR) pathway (37,45,46), their reduction is consistent with a shift toward a more immune-permissive microenvironment. Importantly, spatial “enrichment” within a given core can coexist with overall low pathway abundance at the group level, highlighting the value of spatial analysis to distinguish within-tissue heterogeneity from between-group differences. The interplay between immune phenotypes and local metabolite landscapes offers a new dimension for understanding immune resistance mechanisms and refining therapeutic stratification in MuM.

Taken together, these findings suggest that the recruitment of functional immune cell types to tumor regions, a stromal context linked to immune exclusion, and compartment-specific immunometabolic states collectively define microenvironments associated with ICI responsiveness in MuM. The integration of multi-omic spatial profiling offers promise for guiding personalized therapeutic strategies, advancing precision medicine, and ultimately improving clinical outcomes in MuM. Future studies should validate these spatial features in larger tissue regions and independent cohorts, and explore therapeutic interventions targeting stromal barriers and myeloid-metabolic programs to enhance immunotherapy efficacy.

### Limitations of the study

This study has several limitations. First, although the cohort is notable for this rare disease, the sample size remains modest and analyses were performed on FFPE tissue cores, which provide limited spatial coverage and may be sensitive to regional sampling and intratumoral heterogeneity. Second, while COMET enables single-cell spatial mapping of cell phenotypes and neighborhood architecture, the fixed ∼40-plex protein panel necessarily limits the breadth of functional state and ligand–receptor pathway interrogation. Third, MALDI-IMS detects metabolite features with varying confidence of annotation; orthogonal validation of key metabolite identities and pathway activity will strengthen biological interpretation. Finally, the absence of matched genomic and/or transcriptomic profiling in this study precluded direct linkage of oncogenic alterations to spatial TME organization and metabolic reprogramming.

## Methods

### Patient cohorts

The melanoma databases were queried to identify patients with primary MuM that had undergone treatment with immunotherapy targeting the PD1-PD-L1 axis or CTLA4, specifically using Pembrolizumab, Nivolumab and Ipilimumab as combination or monotherapy. The H&E sections of pre-treatment and post-treatment cases were evaluated and paraffin blocks of melanoma before and after therapy with sufficient tissue were selected for the study. Based on tissue availability, we identified more pre-treatment samples compared to post-treatment samples for this study. Two tissue microarrays (TMA) were generated using triplicates of 1.0 mm cores; of each sample, we tried to represent the center and periphery of the tumor, based on tissue availability.

The TMAs contained paired pre- and/or post-treatment samples from 26 patients. After quality control for core integrity and tissue availability, 97 cores were available in TMAs, including 75 pre-treatment cores and 22 post-treatment cores (**Supp. Fig. 3**). The cohort comprised of 15 male and 11 female patients, with 9 sinonasal, 11 anorectal, 2 conjunctival and 4 female urethral primary MuM; please refer to **Supp. Table 1** for further details. The post-treatment samples, where available were assessed using the International Neoadjuvant Melanoma Consortium (INMC) criteria (**Supp. Table 2**) (47).

### Raw image acquisition, image processing of MALDI-IMS data

TMA sections were dewaxed by submerging in fresh xylene for 2 x 3 minutes each. After allowing to dry, fiducial markers were placed on the slides using a diamond scribe and digital images were acquired using an Epson Perfection V600 flatbed document scanner at 4800 dpi.

The sections were coated with 10 mg/mL 1,5-diaminonaphthalene matrix in 50% acetonitrile using an HTX M5 Robotic Reagent Sprayer over 4 passes with the following parameters: a flow rate of 100 µL/min, a track speed of 1200 mm/min, a track spacing of 3 mm, a CC track pattern, a nozzle temperature of 60°C, a nozzle height of 40 mm, and an N2 pressure of 10 psi.

Mass spectrometry images were acquired using FlexImaging 7.0 on Bruker timsTOF fleX QTOF mass spectrometer in negative ion mode with TIMS OFF at 20 µm resolution. A total of 200 laser shots were summed per pixel. The following instrument parameters were used: an m/z range of 50–600, a Funnel 1 RF of 150.0 Vpp, a Funnel 2 RF of 200.0 Vpp, a Multipole RF of 200.0 Vpp, a Collision Energy of 10.0 eV, a Collision RF of 500.0 Vpp, a Transfer Time of 55.0 µs, and a PrePulse Storage of 5.0 µs.

Data were imported into SCiLS Lab 2025b (Bruker) and root mean square normalized. Metabolite peaks were manually chosen with an integration window of ±15 ppm. Peaks corresponding to MALDI matrix and non-monoisotopic peaks were avoided. Putative metabolite IDs were assigned using the MetaboScape (Bruker) plugin for SCiLS lab, searching against the HMDB and other vendor provided databases, allowing an m/z tolerance of 10 ppm. Negative mode ions of [M-H]-, [M]-, and [M+Cl]- were considered. Images of selected identified metabolites were exported to OME.tiff files for further analysis.

### Sequential immunofluorescence using the COMET platform

Formalin-fixed, paraffin-embedded tissue microarray sections (5 μm) were baked overnight at 60°C. Slides were subsequently dewaxed and rehydrated, followed by melanin bleaching using 0.5% hydrogen peroxide (H₂O₂) in Tris-HCl buffer (pH 10; Sigma-Aldrich, St. Louis, MO, USA) at 80°C for 15 minutes. Heat-induced antigen retrieval was then performed in Tris-EDTA buffer (pH 9.0; BioGenex, HK549-XAK) at 97°C for 15 minutes using the EZ-Retriever system (BioGenex, MW014-MO). After allowing the slides to cool to room temperature for 30 minutes, they were washed three times with Multi-staining Buffer (MSB, Lunaphore BU06; 1:20 in deionized water) and stored hydrated in MSB at room temperature until processing (48).

Multiplex immunofluorescence staining was performed on the Lunaphore COMET NSP system, following the manufacturer’s sequential immunofluorescence (seqIF) protocol (49). Staining was carried out in automated cycles, each consisting of incubation with off-the-shelf primary antibodies (two markers per cycle; see **Supp. Table 3**), followed by Alexa Fluor™ 555 or 647-conjugated secondary antibodies. Imaging was performed after each cycle using the integrated widefield microscope (20×/0.7 NA objective; DAPI, TRITC, Cy5 channels). Antibodies were gently eluted between cycles using proprietary Lunaphore reagents.

### Image preprocessing and alignment

The resulting OME-TIFF image stacks (0.23 μm/pixel resolution) were processed with flat-field correction and autofluorescence subtraction to mitigate uneven illumination and background signals. Image alignment between the MALDI-IMS and COMET datasets, as well as tissue and cell segmentation followed by spatial analyses, were performed using Visiopharm software (VIS, version 2024.07.1.16912; Visiopharm, Denmark).

Serial tissue sections were co-registered via the Tissuealign module employing automated tissue outline matching, further refined manually by placement of landmarks on corresponding morphological features as necessary. Quality control was ensured through overlay visualization and quantitative assessment of registration accuracy.

### Tissue segmentation and region of interest (ROI) definition

Tumor ROIs were delineated based on SOX10 expression using Visiopharm’s machine learning tissue classification module. The remaining ROIs within tissue cores were defined as TME ROIs. Classification models were trained on representative pathologist-annotated images and subsequently reviewed and confirmed by board-certified pathologists to validate segmentation accuracy.

### Cell segmentation and phenotyping

Cell segmentation was conducted using Visiopharm’s integrated deep learning nuclei detection algorithm. Nuclei were expanded to approximate whole-cell segmentation by integrating signals from nuclear and cytoplasmic markers. Manual cell phenotyping was performed post-segmentation according to marker criteria outlined in **Supp. Fig. 4**. All segmentation and phenotyping data were exported for downstream quantitative and spatial analyses.

### CN definition

To identify CNs, a window was defined according to the index cell and its surrounding cells within a radius of 70 μm (31). The index cell and its surrounding cells in a window were counted and annotated and further transformed into a proportion for further K-means clustering with k = 15 and allocated to each CN. To validate the CN assignment, these allocations were overlaid on the original tissue H&E-stained and fluorescent images. The comprehensive spatial analysis of cellar neighborhoods were further managed using the SPACEc pipeline (50).

### Patch proximity analysis

Patches were detected in COMET data by running HDBSCAN clustering over all centroids in the specified CN. The resulting clusters were used as patches, and cells that were not part of a patch are ignored. The outermost cells of each patch were selected by constructing a concave hull and selecting the spanning points. To include cells close to the outlining border of the patch and to correct for cells that were distant from the patch, three nearest neighbors for each edge cell were included as well, resulting in a group of cells surrounding each patch. Subsequently, the selected cells were used as anchor cells to span a radius (5μm, 10μm, 15μm, 20μm, and 25μm) around each selected cell. Cells within the radius that did not belong to the patch were counted as cells within spatial proximity.

### Cell-cell interaction analysis

Briefly, Delaunay triangulation for each cell was determined based on x, y positions within each field of view using the default settings from the SciPy package. To identify interacting cells and their coordinates, relevant information was extracted automatically from the output. Subsequently, the distances between cells connected by the edges of a Delaunay triangle in the two-dimensional space were computed using the following formula:

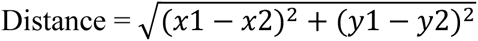

Cell-to-cell interactions within 265 pixels were identified. To establish a reference distribution of distances, 100 iterations of the triangulation calculation were conducted. In each iteration, the cells and their neighborhoods within each field of view were randomly reassigned to existing x, y positions. The average distances of cell-cell interactions for each field of view in each permutation were calculated and compared to the observed distances using a Mann-Whitney U Test. The fold enrichments of distances between the observed data compared to the mean distances derived from the permutation test were determined. The top six values of log fold changes of distance for each pair of interactions with p-values were less than 0.05 were plotted.

### Spatial context map

First, a window size of 70 nearest neighbors was used to create the composition vectors. Within each window, the fewest neighborhoods that made up more than 85% of the neighborhoods was identified. This combination informed about prominent associations of neighborhoods in the window, which was a feature termed spatial context. Third, we counted each combination and connected the most prevalent combinations into a spatial context map. The hierarchical spatial context map showed different levels of neighborhood combinations and their relative frequencies.

### Cellular neighborhood interface analysis

A barycentric coordinate projection was generated by applying window size of 70 nearest neighbors to every cell. Window composition was analyzed in terms of the percentage distribution of CNs. Windows containing less than 80% of the three selected neighborhoods were excluded from the analysis.

### Statistical analyses

Correlations of proteins and metabolites were calculated using pairwise Spearman’s rank-order correlation and p-values were adjusted with Benjamini–Hochberg correction. Volcano plot was used for testing metabolite intensity differences with a cutoff p-value of <0.05 and a fold change of >1. Metabolite enriched pathway analysis was performed via the Kyoto Encyclopedia of Genes and Genomes (KEGG) database (Fisher’s exact test, q < 0.05 for FDR correction). Cell type-based CNs were clustered by K-means clustering analysis for stratify R and NR patients and visualized as heatmap. The Mann–Whitney *U* test was used for testing cell type proportion differences. The Benjamini-Hochberg’s method was used for multiple test correction, and false discovery rate (FDR) was subsequently calculated. The cell density plot was generated using the function plot_3d_density in the software Spyrrow (https://github.com/liuyunho/Spyrrow). The cell patch plot was generated using the cell boundary coordinates, implemented with the function plot_cell_patch in the software Spyrrow, where each cell patch was treated as a polygon object and generated using the Polygon function. Results were considered statistically significant when p-values or FDRs were less than 0.05. All statistical tests were conducted using python.

## Data availability

The data generated in this study are available upon reasonable request from the corresponding author.

## Acknowledgements

We are grateful to the patients for making available the tumor samples that contribute to this research. This work was supported by The University of Texas, MD Anderson Cancer SPORE in Melanoma, P50-CA093459 (**S. Ekmekcioglu**), Melanoma Research Alliance, #570806 (**P. Nagarajan**), MD Anderson Cancer Center Support Grant, P30-CA016672 (**S. Ekmekcioglu and J. Burks**), NCI Research Specialist 1 R50 CA243707-01A1 (**J. Burks**), Shared Instrumentation Award from the Cancer Prevention Research Institution of Texas (CPRIT), RP121010 (**J. Burks**), Foundation for the National Institutes of Health (FNIH)-PACT (**S. Ekmekcioglu**), Scientific and financial support for the PACT project are made possible through funding support provided to the FNIH by AbbVie Inc., Amgen Inc., Boehringer-Ingelheim Pharma GmbH & Co. KG, Bristol-Myers Squibb, Celgene Corporation, Genentech Inc., Gilead, GlaxoSmithKline plc, Janssen Pharmaceutical Companies of Johnson & Johnson, Novartis Institutes for Biomedical Research, Pfizer Inc., and Sanofi. **L.Wang** was in part supported by the James P. Allison Institute and the Institute for Data Science in Oncology at the University of Texas MD Anderson Cancer Center.

## Author contributions

**J. Wang:** Data curation, formal analysis, data interpretation, investigation, visualization, methodology, writing–original draft, writing–review and editing. **P. Nagarajan:** Resources, funding acquisition, investigation, writing–review and editing. **S. Cho:** Data curation, experimental work, methodology. **Yunhe Liu:** Methodology, visualization, data interpretation, writing–review and editing. **E. Seeley**: Methodology. **Y. Dai, Yang Liu and K. Yu:** Data interpretation**. J. Burks, J. McQuade and A. Diab:** Resources. **L. Wang:** Conceptualization, supervision, visualization, data interpretation, writing–original draft, writing–review and editing. **S. Ekmekcioglu:** Conceptualization, supervision, funding acquisition, project administration, writing–original draft, writing–review and editing.

## Notes

**Conflict of interest statement:** L.Wang serves as a member of the Scientific Advisory Board for SELLAS Life Sciences and receives compensation outside the scope of this submitted work. All other authors declare no competing interest.

### Competing Interest Statement

L.Wang serves as a member of the Scientific Advisory Board for SELLAS Life Sciences and receives compensation outside the scope of this submitted work. All other authors declare no competing interest.

