## Supplemental Table 1 for "Integration of spatial single-cell proteomics and spatial metabolomics reveals tumor microenvironment predictive of immunotherapy response in mucosal melanoma"

Supplementary Table 1. Cohort characteristics.

|  |  | | Total |
| --- | --- | --- | --- |
| Number | All cores | | 97 |
|  | Pre-treatment cores | | 75 |
|  | Post-treatment cores | | 22 |
|  | Patients | | 26 |
|  | Male patients | | 15 |
|  | Female patients | | 11 |
| Primary mucosal melanoma | Sinonasal | | 9 |
|  | Anorectal | | 11 |
|  | Conjunctival | | 2 |
|  | Female genitourinary | | 4 |
| Core location within tumor | Periphery | | 50 |
|  | Center | | 47 |
| Core tissue composition | Tumor-enriched | | 46 |
|  | Interface-enriched | | 35 |
|  | Stroma-enriched | | 16 |
| Density of inflammatory infiltrate | Sparse | | 44 |
|  | Mild | | 14 |
|  | Moderate | | 34 |
|  | Dense | | 5 |
| INMC pathological tumor response to therapy | Non-responder (NR) | No response, NR (tumor viability >50%) | 20 |
|  | Responder (R) | Partial response, PR (tumor viability >10 to ≤50%) | 3 |
|  |  | Major response, MR (tumor viability 1 to ≤10%) | 1 |
|  |  | Complete response, CR (tumor viability 0%) | 2 |
