## Supplemental Table 2 for "Integration of spatial single-cell proteomics and spatial metabolomics reveals tumor microenvironment predictive of immunotherapy response in mucosal melanoma"

Supplementary Table 2. Tumor response to immunotherapy, by International Neoadjuvant Melanoma Consortium (INMC) criteria.

|  | Tumor bed composition | | | | Pathological response by INMC criteria |
| --- | --- | --- | --- | --- | --- |
| Patient number | % Viable tumor | % Necrosis | % Fibrosis | % Tumoral melanosis |  |
| P1 | 100 | 0 | 0 | 0 | NR |
| P2 | 80 | 5 | 5 | 10 | NR |
| P3 | 90 | 3 | 0 | 7 | NR |
| P4 | 100 | 0 | 0 | 0 | NR |
| P5 | 0 | 30 | 10 | 60 | CR |
| P6 | 10 | 10 | 60 | 20 | MR |
| P7 | 100 | 0 | 0 | 0 | NR |
| P8 | 100 | 0 | 0 | 0 | NR |
| P9 | 40 | 0 | 50 | 10 | PR |
| P10 | 100 | 0 | 0 | 0 | NR |
| P11 | 95 | 0 | 0 | 5 | NR |
| P12 | 80 | 18 | 0 | 2 | NR |
| P13 | 17 | 0 | 79 | 4 | PR |
| P14 | 70 | 2 | 15 | 3 | NR |
| P15* | NA | NA | NA | NA | NR |
| P16* | NA | NA | NA | NA | NR |
| P17 | 90 | 0 | 10 | 0 | NR |
| P18 | 100 | 0 | 0 | 0 | NR |
| P19 | 95 | 5 | 0 | 0 | NR |
| P20 | 20 | 80 | 0 | 0 | PR |
| P21 | 100 | 0 | 0 | 0 | NR |
| P22 | 100 | 0 | 0 | 0 | NR |
| P23 | 60 | 0 | 30 | 10 | NR |
| P24 | 0 | 0 | 100 | 0 | CR |
| P25 | 95 | 0 | 0 | 5 | NR |
| P26 | 95 | 0 | 0 | 5 | NR |

* Two patients, P16 and P17 showed clinical progression and did not receive post-treatment surgical excision and thus, were considered to be non-responders (NR).

INMC criteria for pathological response: no response, NR (tumor viability >50%); partial response, PR (tumor viability >10 to ≤50%); major response, MR (tumor viability 1 to ≤10%); complete response, CR (tumor viability 0%)
