## Supplemental Table 3 for "Integration of spatial single-cell proteomics and spatial metabolomics reveals tumor microenvironment predictive of immunotherapy response in mucosal melanoma"

Supplementary Table 3. Antibody Panel Utilized for COMET Multiplex Immunofluorescence

| Cycle | Target | Clone | Host | Vendor | Catalog | Dilution | Channel |
| --- | --- | --- | --- | --- | --- | --- | --- |
| 1 | RORgt | 6F3.1 | Mouse | FortisLife | ACI3208B | 250 | TRITC |
| 1 | CD25 (IL2RA) | EPR6452 | Rabbit | Abcam | Ab128955 | 50 | Cy5 |
| 2 | FOXP3 | 236A/E7 | Mouse | eBioscience | 14-4777-82 | 50 | TRITC |
| 2 | CD3e | BL-298-5D | Rabbit | FortisLife | A700-016CF | 400 | Cy5 |
| 3 | Granzyme B | GrB-7 | Mouse | Agilent Dako | M723501-2 | 50 | TRITC |
| 3 | CD123 (IL3RA) | Polyclonal | Rabbit | Millipore-Sigma | HPA003539 | 100 | Cy5 |
| 4 | CD163 | 10D6 | Mouse | Invitrogen | MA5-1145 | 50 | TRITC |
| 4 | IRF4 | Polyclonal | Rabbit | Millipore Sigma | HPA002038 | 100 | Cy5 |
| 5 | CD303 (BDCA2) | 992258 | Mouse | R&D | AMAB62991 | 100 | TRITC |
| 5 | Tbet | BLR110H | Rabbit | Bethyl | A700-110 | 100 | Cy5 |
| 6 | CD31 | 3F8E2 | Mouse | Proteintech | 66065-2-Ig | 250 | TRITC |
| 6 | Arginase-1 | D4E3M | Rabbit | CST | 93668 | 100 | Cy5 |
| 6 | Arginase-1 | BLR036F | Rabbit | FortisLife | A700-036 | 100 | Cy5 |
| 7 | HLA-DR | LN3 | Mouse | FortisLife | A500-022A | 100 | TRITC |
| 7 | LAG3 | Polyclonal | Rabbit | Proteintech | 16616-1-A | 100 | Cy5 |
| 8 | CD74 | PIN.1 | Mouse | Novus bio | NB100-198 | 100 | TRITC |
| 8 | IRF8 | polyclonal | Rabbit | Millipore Sigma | HPA002267 | 100 | Cy5 |
| 9 | CD68 | KP-1 | Mouse | FortisLife | A500-018AF | 250 | TRITC |
| 9 | CXCR4 | D4Z7W | Rabbit | CST | 90927 | 100 | Cy5 |
| 10 | CD208 (Lamp3) | 1010E1.01 | Rat | Novus Bio | DDX0191P-100 | 100 | TRITC |
| 10 | iNOS | polyclonal | Rabbit | Novus bio | NB300-605 | 100 | Cy5 |
| 11 | CD8a | C8/144B | Mouse | FortisLife | A500-021AF | 100 | TRITC |
| 11 | CD86 | E2G8P | Rabbit | CST | 76755SF | 100 | Cy5 |
| 12 | CD20 | L26 | Mouse | FortisLife | A500-017AF | 250 | TRITC |
| 12 | CD4 | EPR6855 | Rabbit | Abcam | ab181724 | 100 | Cy5 |
| 13 | CD141 | 15C8 | Mouse | Leica Biosystems | CD141-L-U | 100 | TRITC |
| 13 | CD206 | BLR109H | Rabbit | FortisLife | A700-109 | 500 | Cy5 |
| 14 | CD1c | 2A7C11? | Mouse | Novus Bio | NBP2-61726 | 100 | TRITC |
| 14 | CD56 | BLR152J | Rabbit | FortisLife | A700-152 | 1000 | Cy5 |
| 15 | CD45RO | UCHL1 | Mouse | CST | 36282 | 1000 | TRITC |
| 15 | CD11b | EPR1344 | Rabbit | AbCam | ab133357 | 500 | Cy5 |
| 16 | CD45RA | IHC-536-C | Mouse | GenomeMe | IHC-536-C | 200 | TRITC |
| 16 | Sox10 | EPR4007-104 | Rabbit | Abcam | ab180862 | 100 | Cy5 |
| 17 | MIF | polyclonal | Goat | R&D | AF289-PB | 200 | TRITC |
| 17 | CD14 | EPR3635 | Rabbit | Millipore-Sigma | 114R-16 | 500 | Cy5 |
| 18 | CD44 | 156-3C11 | Mouse | Abcam | ab213072 | 500 | TRITC |
| 18 | Ki-67 | EPR3610 | Rabbit | Abcam | ab209897 | 200 | Cy5 |
| 19 | a-SMA | 1A4 | Mouse | CST | 69319 | 100 | TRITC |
| 19 | CD11c | EP1347Y | Rabbit | Abcam | ab216655 | 1000 | Cy5 |
| 20 | Vimentin | IHC-684-C | Mouse | GenomeMe | IHC-684-C | 1000 | TRITC |
| 20 | CD45 | BL-178-12 | Rabbit | FortisLife | A700-012 | 1000 | Cy5 |
