## Supplementary figures and images for "Integration of spatial single-cell proteomics and spatial metabolomics reveals tumor microenvironment predictive of immunotherapy response in mucosal melanoma"

### Supplemental figure 1

Supplementary fig1

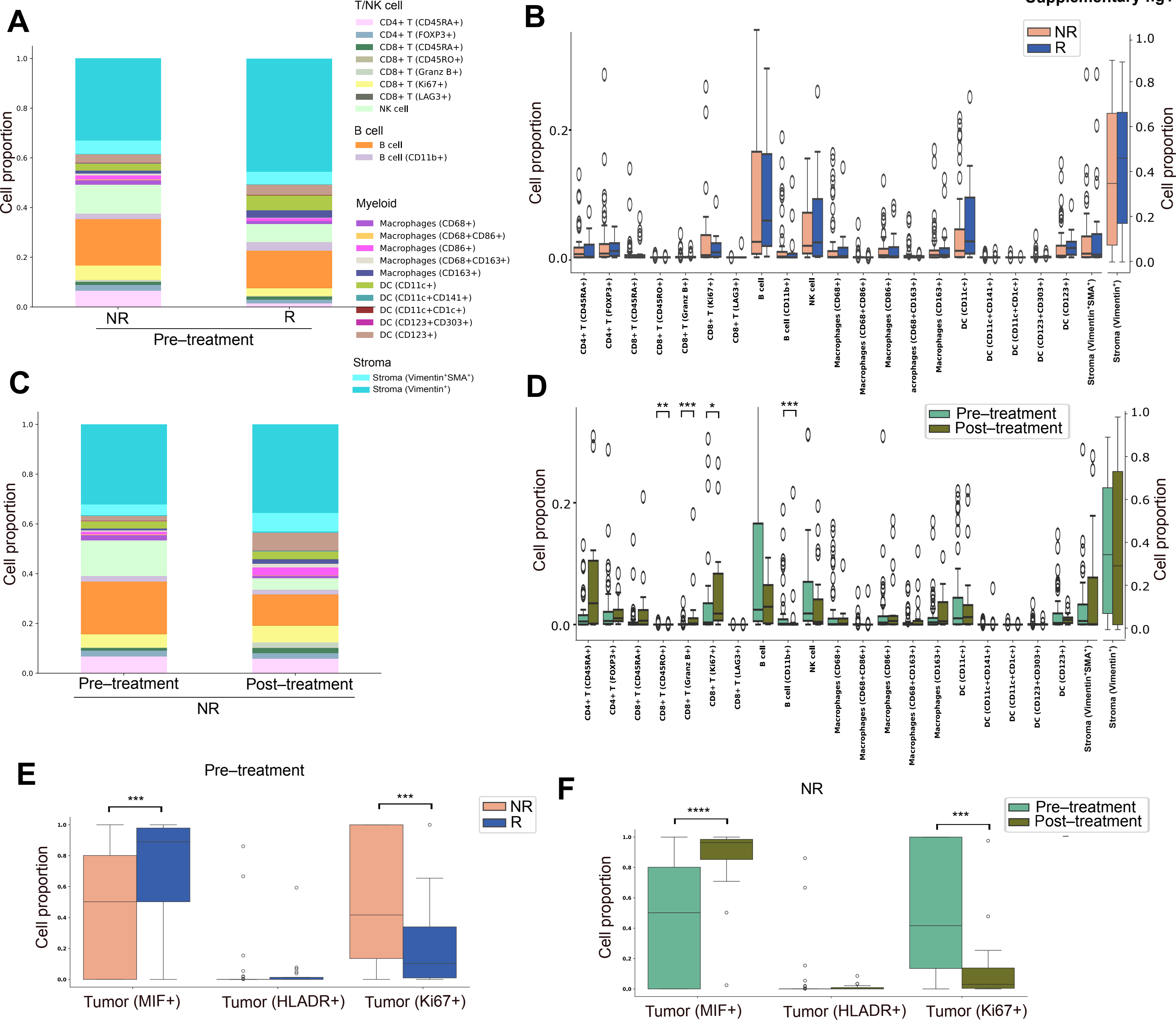

### Supplemental figure 2

#### Supplementary fig2

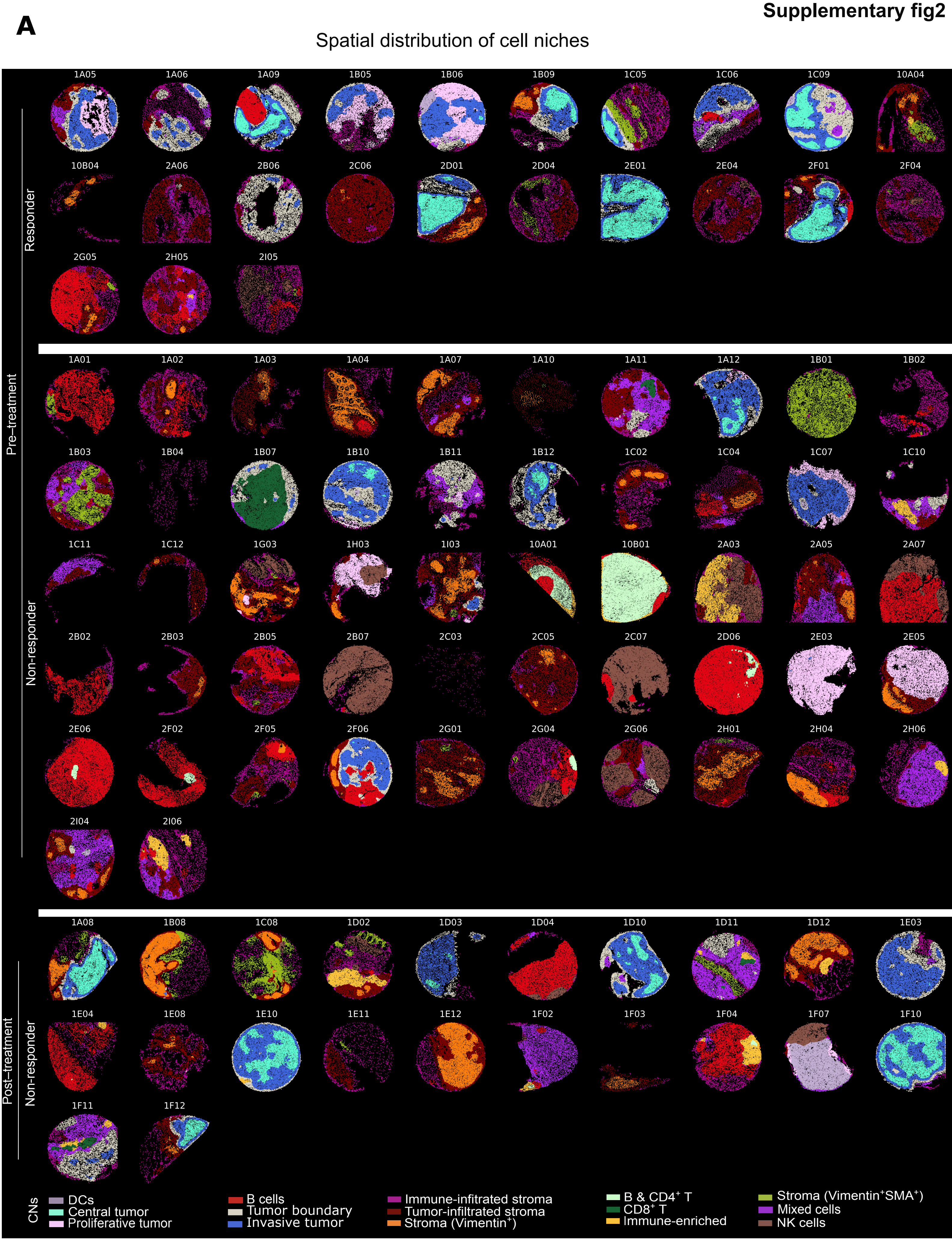

### Supplemental figure 3

**Supplementary fig3**

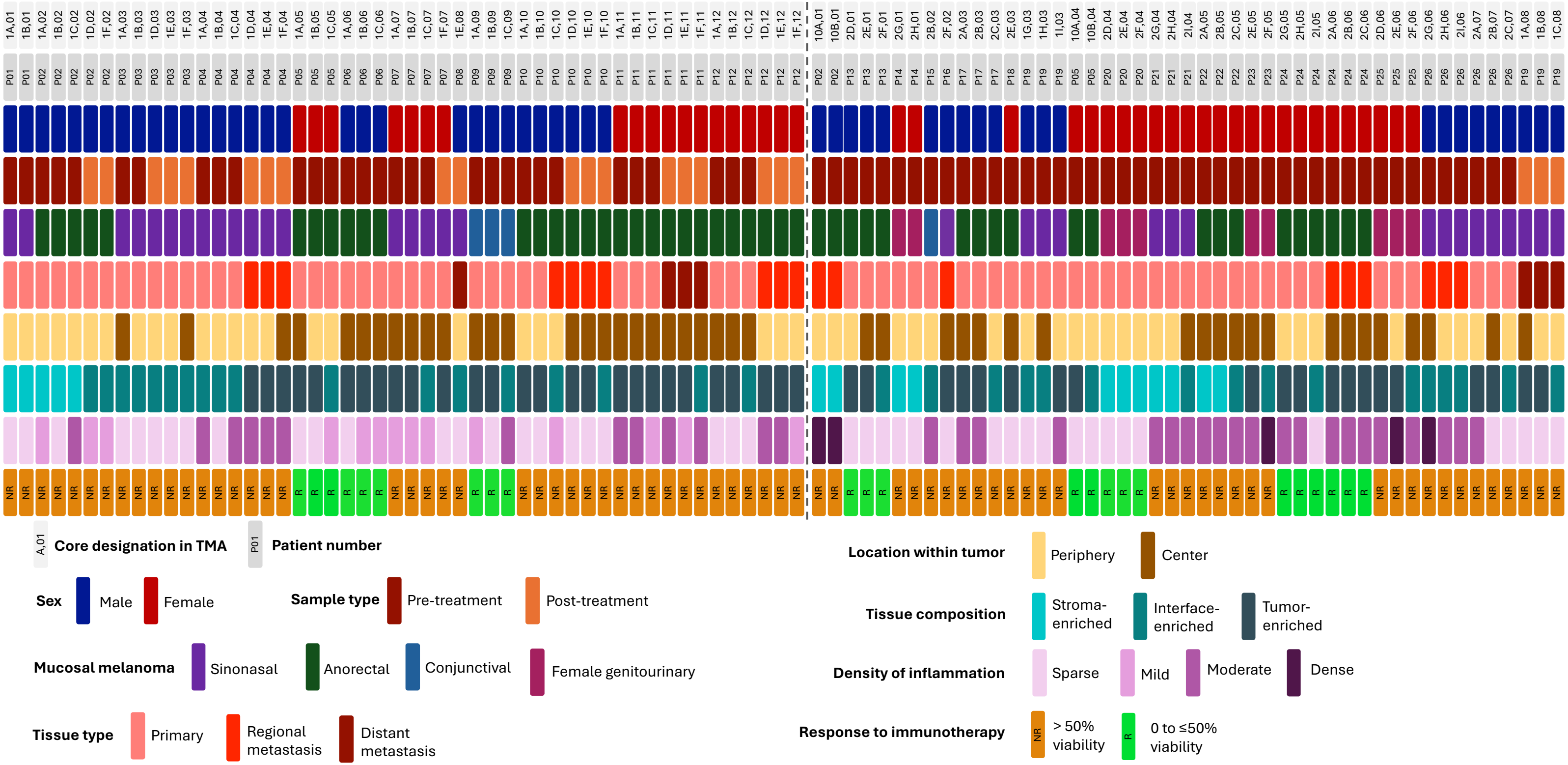

### Supplemental figure 4

**Supplementary fig4**

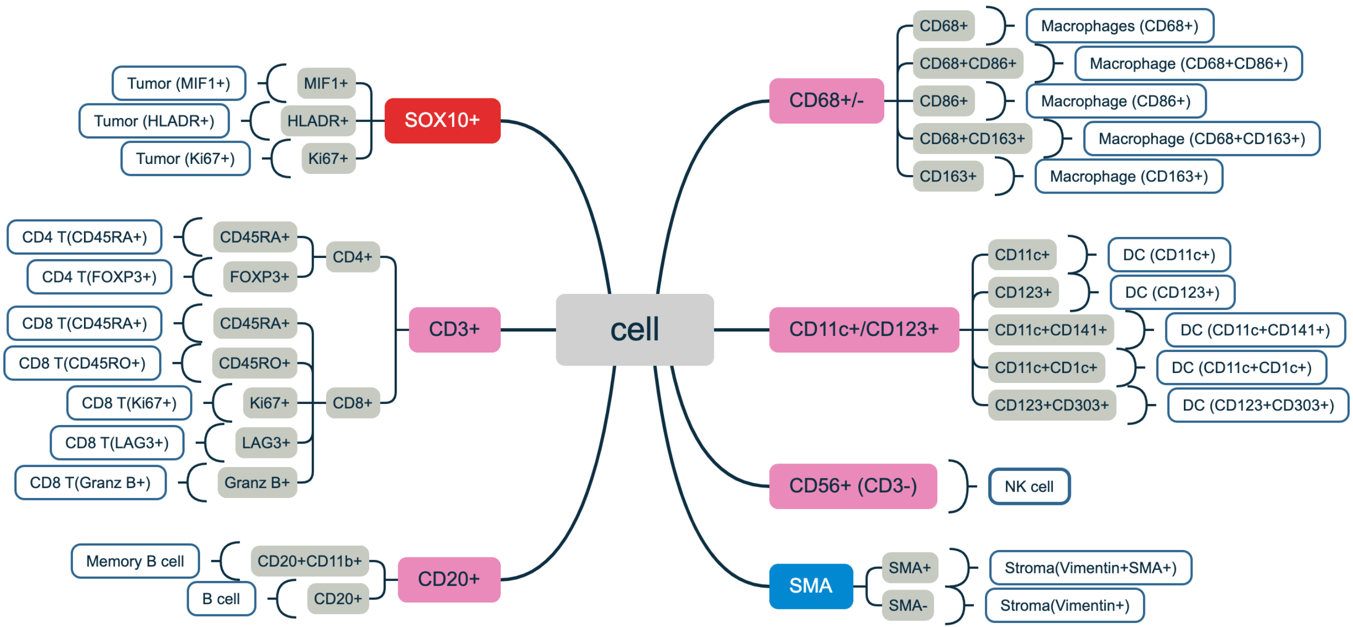
